# CNS Glycosylphosphatidylinositol Deficiency Results in Delayed White Matter Development, Ataxia, and Premature Death in a Novel Mouse Model

**DOI:** 10.1101/763490

**Authors:** Marshall Lukacs, Rolf W. Stottmann

## Abstract

The Glycosylphosphatidylinositol (GPI) anchor is a post-translational modification added to approximately 150 different proteins to facilitate proper membrane anchoring and trafficking to lipid rafts. Biosynthesis and remodeling of the GPI anchor requires the activity of over twenty distinct genes. Defects in the biosynthesis of GPI anchors in humans leads to Inherited Glycosylphosphatidylinositol Deficiency (IGD). IGD patients display a wide range of phenotypes though the central nervous system (CNS) appears to be the most commonly affected tissue. A full understanding of the etiology of these phenotypes has been hampered by the lack of animal models due to embryonic lethality of GPI biosynthesis gene null mutants. Here we model IGD by genetically ablating GPI production in the CNS with a conditional mouse allele of *phosphatidylinositol glycan anchor biosynthesis, class A (Piga)* and *Nestin-Cre.* We find that the mutants do not have structural brain defects but do not survive past weaning. The mutants show progressive decline with severe ataxia consistent with defects in cerebellar development. We show the mutants have reduced myelination and defective Purkinje cell development. Surprisingly we found *Piga* was expressed in a fairly restricted pattern in the early postnatal brain consistent with the defects we observed in our model. Thus, we have generated a novel mouse model of the neurological defects of IGD which demonstrates a critical role for GPI biosynthesis in cerebellar and white matter development.

## Introduction

Inherited Glycosylphosphatidylinositol Deficiency (IGD) is defined by a deficiency of the cell surface glycosylphosphatidylinositol (GPI) anchor [1, 2]. The GPI anchor is a glycolipid added post-translationally to nearly 150 proteins and is required for membrane anchoring and lipid raft trafficking of these proteins [1, 3]. Over twenty genes are required for the biosynthesis and remodeling of the GPI anchor in the endoplasmic reticulum and Golgi apparatus. IGD patients display a wide array of phenotypes including neural, craniofacial, cardiac, renal, hepatic, ophthalmologic, skeletal, dental, dermatologic, and sensorineural defects [1, 2, 4]. This wide array of phenotypes highlights the broad requirement for GPI biosynthesis and remodeling in the development of many organ systems.

A recent review of IGD patients found that the most commonly affected organ system, regardless of the mutated gene in the GPI biosynthesis pathway, is the central nervous system (CNS) [2]. These CNS phenotypes include structural CNS deficits including microcephaly, hypoplastic corpus callosum, hypoplastic cerebellum, white matter immaturity, and cortical atrophy. Clinical findings related to these structural deficits include epilepsy, developmental delay, intellectual disability, hypotonia, hyperreflexia, chorea, ataxia, and behavioral abnormalities with variable penetrance [2, 5–7]. Electroencephalogram imaging reveals abnormal electrical activity including hypsarrhythmia indicative of abnormal brain development in nearly all patients [2, 8, 9]. Often the epilepsy is intractable and severely impairs quality of life. Some reports have shown Vitamin B_6_ is beneficial in treating IGD-related epilepsy. It has been hypothesized the benefit of Vitamin B_6_ is related to the defects in GPI-anchored tissue nonspecific alkaline phosphatase which dephosphorylates B_6_ thereby allowing it to pass the membrane and participate in in γ-aminobutyric acid (GABA) synthesis [10, 11]. However, it is unclear if all patients respond to Vitamin B6 supplementation and no rigorous study has tested this hypothesis, at least in part due to a relative lack of mouse models of IGD. Little is known about the requirement for GPI biosynthesis in the CNS or the mechanisms that lead to these clinical phenotypes. Therapy is currently limited to symptomatic treatment and most patients will die before age 5 by cardiac arrest, aspiration pneumonia, or central respiratory failure [12]. Thus, there is a clear need to understand the pathophysiology that results from GPI deficiency in the CNS.

The initiation of GPI biosynthesis requires the Phosphatidylinositol Glycan Anchor Biosynthesis Class A (PIGA) protein, a component of the phosphatidylinositol N-acetylglucosaminyltransferase complex. This complex generates N-acetylglucosamine-phosphatidylinositol (GlcNac-PI) by transferring N-acetylglucosamine (GlcNac) from UDP-GlcNac to phosphatidylinositol (PI). GlcNac-PI is the first precursor in the biosynthesis of the GPI anchor and deletion of *Piga* completely abolishes GPI biosynthesis [1, 3]. Patients with germline mutations in *PIGA* develop a form of IGD termed Multiple Congenital Anomalies-Hypotonia-Seizures Syndrome 2 (MCAHS2: OMIM 300868) [13]. A recent review of 10 MCAHS2 patients found all had seizures, abnormal EEG findings, developmental delay and intellectual disability. 8/10 patients had hypotonia and 5/10 had hyperreflexia, cerebellar hypoplasia, and cortical atrophy [12]. 7/10 patients developed white matter defects including thin corpus callosum and white matter immaturity. Thus, *PIGA* deficiency serves as a representative model to study the broad requirement for GPI biosynthesis in CNS development.

Other groups have taken a genetic approach to ablate GPI biosynthesis in specific mouse tissues by combining a conditional allele of *Piga* with flanking loxP sites (*Piga^flox^*) with tissue specific Cre recombinase transgenics. Recently, we used this conditional allele to delete *Piga* in the developing mouse neural crest cells with *Wnt1-Cre.* This deletion resulted in cleft lip and cleft palate as well as hypoplasia of the craniofacial skeleton highlighting a critical role for GPI biosynthesis in neural crest cell survival [14]. Others have used this approach to reveal an important role for GPI biosynthesis in skin, limb, blood and craniofacial development [15–17]. However, germline knockout of *Piga* using CMV-Cre resulted in embryonic lethality with neural tube defect, edema, and cleft lip/palate precluding the analysis of any postnatal neurological phenotype [18]. To overcome this, we sought to conditionally ablate GPI biosynthesis after the closure of the neural tube, at embryonic day E9.5 in mouse. To completely abolish CNS GPI biosynthesis at this stage, we chose to use *Nestin-Cre* to drive deletion of *Piga* broadly in the central and peripheral nervous system starting at E11.5 [19]. As *Piga* resides on the X-chromosome, we generated *Piga^flox/Y^*, Nes-Cre+ hemizygous conditional knockout males (hemizygous cKO) with complete deletion of *Piga* in the *Nestin* lineage and *Piga^flox/WT^, Nes-Cre* mosaic cKO females (mosaic cKO) containing one conditional allele of *Piga* and one wild-type *Piga* allele. Due to random X inactivation, these females develop a mosaic cKO of *Piga* in the *Nestin* lineage. Both hemizygous cKO and mosaic cKO animals survived to postnatal stages and developed some of the CNS phenotypes observed in IGD patients including white matter immaturity, gait imbalance, motor incoordination, and early death. Our conditional knockout approach allowed us to investigate the requirement for GPI biosynthesis for the first time *in vivo* in the mammalian CNS. This model may also be used to test novel IGD therapeutic interventions in the future.

## Results

### *Piga* is expressed broadly in the CNS with enrichment in the corpus callosum and Purkinje cells

We initially sought to determine the expression of *Piga* in the embryonic and postnatal mouse brain. Published and publicly available single cell RNA sequencing data sets are now a robust tool to query gene expression. Zhang and colleagues performed one such experiment in the P7 mouse cortex and found *Piga* is broadly expressed in a variety of neural cell types, but with a clear enrichment in astrocytes and newly formed and myelinating oligodendrocytes [20] (Fig. 1A). We further explored *Piga* expression with immunofluorescence in the postnatal CNS with a commercially available polyclonal αPIGA antibody. We found PIGA protein is enriched in the corpus callosum and cortical periventricular areas at P0 (Fig. 1 B,C). At P10, higher expression was maintained in the periventricular areas and PIGA expression was evident in the Purkinje cells of the cerebellum at this stage (Fig. 1D,E). Further examination revealed clear immunoreactivity in both the cytoplasm and dendrites of the Purkinje cell (Fig. 1E). We concluded from these data that PIGA shows increased expression at P0 in developing white matter areas such as the corpus callosum, consistent with the P7 RNA-Seq data with highest expression in myelinating oligodendrocytes. Once the mature morphology of the cerebellum is established at P10, PIGA is strongly expressed in the Purkinje cells, the sole output of all motor coordination in the cerebellar cortex.

**Figure 1.**
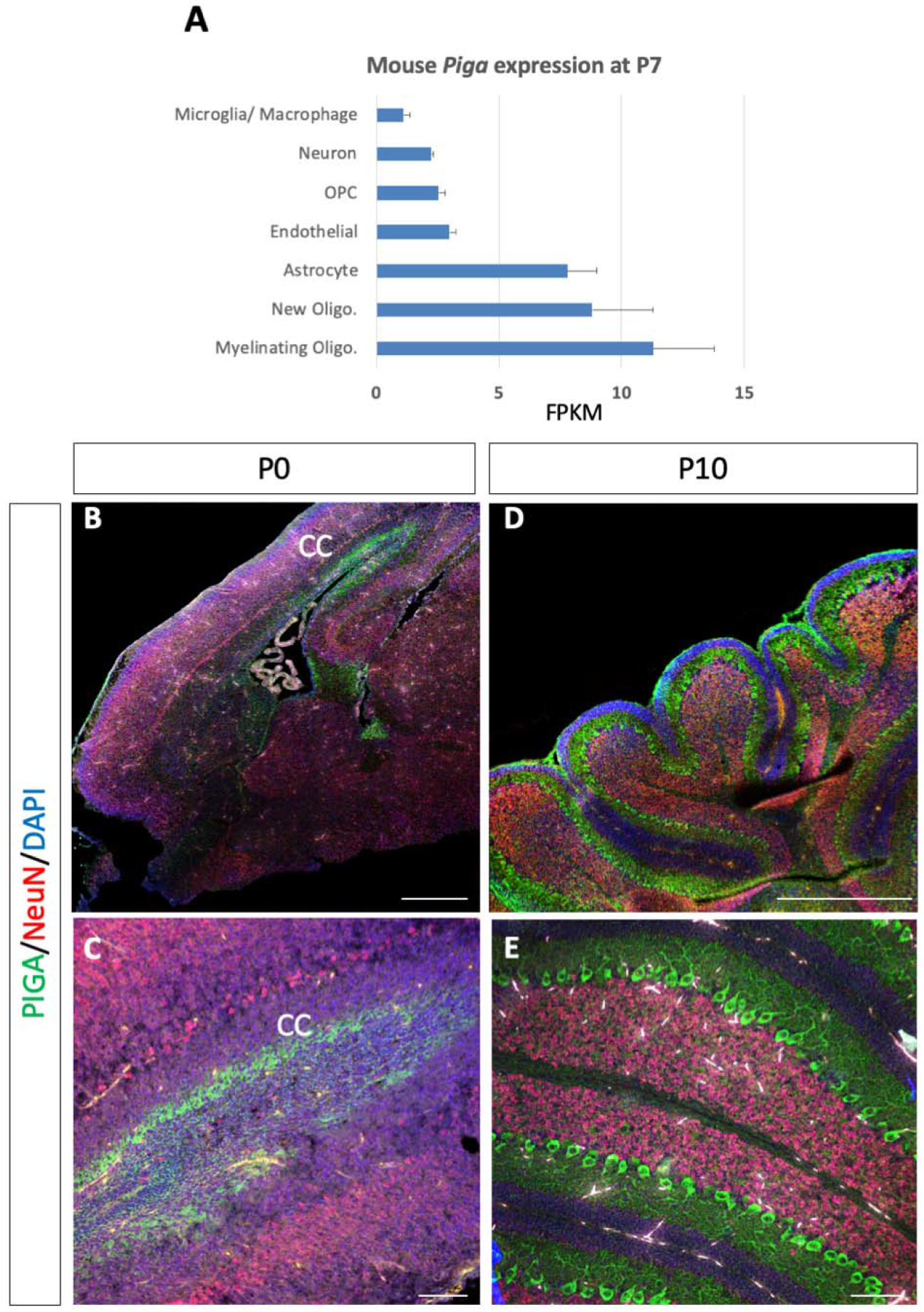
*Piga* expression in the CNS. Data in A is replotted directly from [20] (brainrnaseq.org). P0 PIGA (Green) expression in the corpus callosum (CC) and periventricular areas (B,C). P10 PIGA expression in the cerebellar Purkinje cells D,E). PIGA is shown as green NeuN as a neuronal marker is in red, DAPI counterstain is blue. Fb; forebrain, Mb; Midbrain, BA1;Branchial Arch 1, MNP; Medial Nasal Process, Lb; Limb bud, Oligo.: Oligodendrocyte, OPC: Oligodendrocyte Progenitor Cell. (Scale bars = 500 μm in B,D, 100 μm C,E)

### Deletion of *Piga* in the *Nestin-Cre* lineage results in CNS GPI deficiency

As very few studies have focused on the role of glycosylation in neural development, we aimed to first take a broad approach and delete GPI biosynthesis in the entire CNS/PNS. We hypothesized this would illuminate the critical requirement for GPI biosynthesis in neural development beyond neurulation. We are unable to study the effect of GPI biosynthesis at later neurogenesis stages in the germline *Piga* null mice because these mutants develop neural tube defects or die very early in development [18].

We obtained the conditional *Piga* allele, (*B6.129-Piga^<tm1>^*) which has been used in several studies to delete *Piga* with Cre/lox technology [14, 16, 18]. Upon Cre recombination, the final exon of the conditional *Piga* is deleted and GPI biosynthesis is severely disrupted indicating the allele efficiently abolishes *Piga* function [14, 16, 18]. We crossed these mice to *Nestin-Cre* mice *(B6.Cg-Tg(Nes-cre)1^Kln/J^*). This *Nestin-Cre* allele has been shown to mediate *loxP* excision at E11.5 in the forebrain, midbrain, hindbrain, and spinal cord and maintain expression in neural stem cells throughout development [19, 21, 22]. Therefore, targeting by this *Nestin-Cre* transgene is broad and includes neurons, astrocytes, and oligodendrocytes.

*Piga* is located on the X chromosome in mouse and human resulting in different degrees of *Piga* deletion *flox* flox/Y in male versus female progeny from *Piga^flox^* crosses with *Nestin-Cre.* Males with one copy of Piga^flox/Y^; Nestin-Cre+^/-^ are termed hemizygous conditional knockout (hemizygous cKO) males and should lack all *Piga* expression in the *Nestin* lineage. Females used in this study are mosaic cKOs with the genotype Piga^flox/X^; *Nestin* Cre^+/-^ carrying one floxed allele and one wild-type allele of *Piga*. We chose to continue our behavioral studies with mosaic cKO females because they survive significantly longer than hemizygous cKO males and develop phenotypes similar to those observed in IGD patients including ataxia, decreased lifespan, and neurological decline.

Mice were analyzed from E16.5-postnatal stages. To confirm the Nestin-Cre transgenic was ablating *Piga* in the CNS, we performed PCR with three primers spanning the targeted region to differentiate between wild-type, floxed *(Piga flox)* and recombined *(Piga del)_alleles* (Fig. 2A,B). We found *Nestin-Cre* targeted *Piga* in multiple CNS tissues in *Piga* hemizygous cKO mice as evidenced by the *Piga* deleted PCR product in hemizygous cKO males. Tissues demonstrating *Piga* deletion included the cortex, cerebellum, subcortical structures including the thalamus, hypothalamus, midbrain, pons, and medulla (Fig. 2B, lane 2-4). However, *Piga* excision was not noted in genomic DNA from the tail of the same animal, indicating that the targeting was specific and consistent with previous reports using the Nestin-Cre (Fig. 2B, lane 1). We did observe some remaining *Piga* flox PCR products in the CNS tissues indicating that the Cre did not target absolutely every cell in these tissues. We suspect this *Piga* DNA may be derived from non-neural tissues such as blood vessels, blood, and microglia which are not targeted by *Nestin-Cre.* We conclude our conditional approach targets *Piga* for deletion in the CNS.

**Figure 2.**
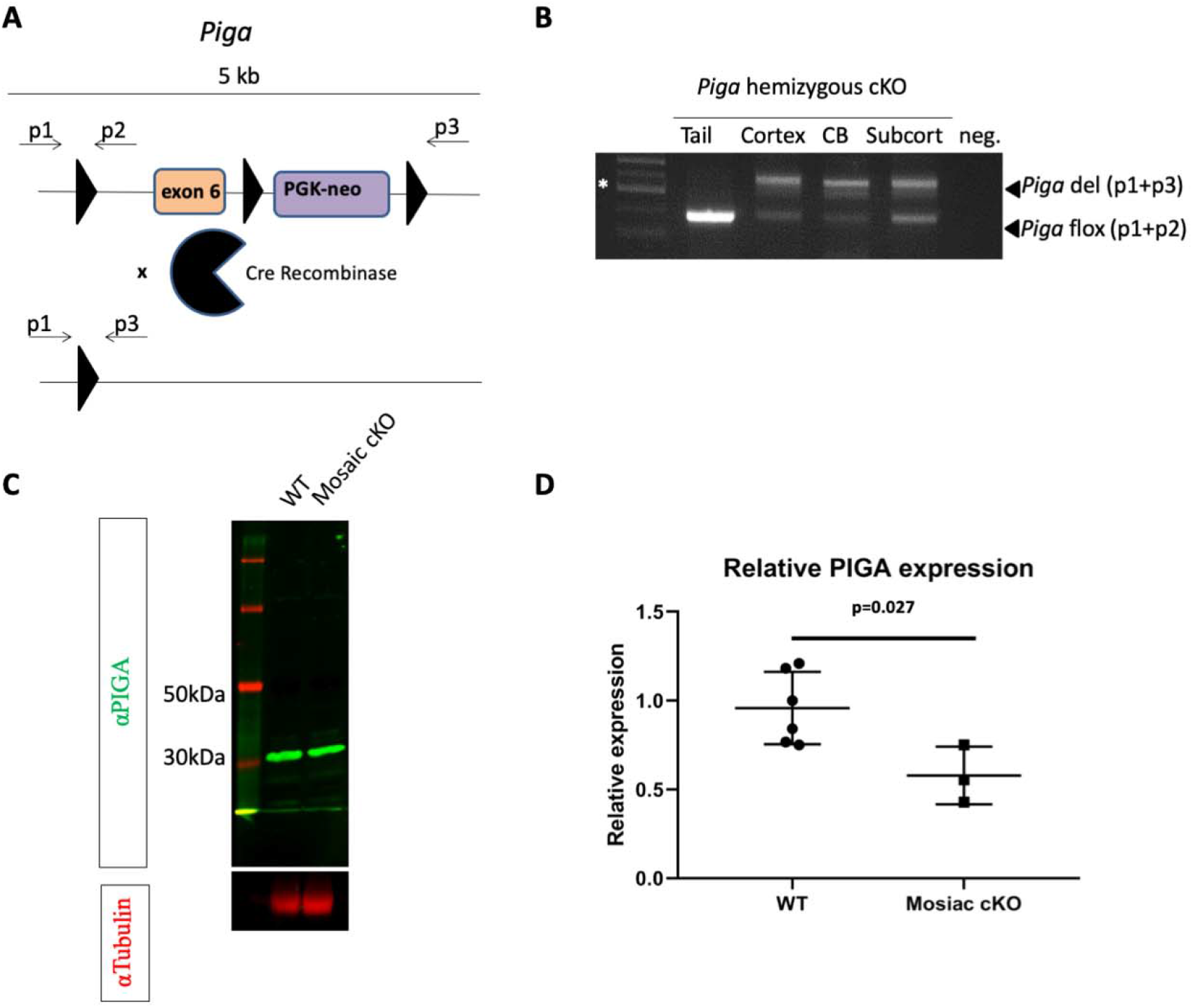
Deletion of *Piga* in the *Nestin-Cre* lineage results in CNS GPI deficiency. Genotyping strategy for identifying *Piga* flox (primers1 and 2 generate a 350bp product), *Piga* deleted (del) (primers 1 and 3 generate a 600bp product) (A). PCR of tail and various CNS structures of Piga hemizygous cKO male mouse (B). Western blot of PIGA (Green) in subcortical tissues from WT and *Piga* Mosaic cKO female littermates, and Tubulin loading control (Red) (C). Quantification of PIGA/tubulin signal from Western blot of N=6 WT and N=3 Mosaic cKO subcortical lysates (D). CB= cerebellum, Subcort=subcortical tissues.

It is known that mothers with one mutant allele of *Piga* display skewing of X inactivation to silence the affected X chromosome carrying the mutant allele [12]. Thus, we were concerned skewing toward the wild-type X chromosome would result in residual PIGA expression higher than the expected 50% of wild-type in mosaic cKOs. To confirm the PIGA protein was reduced in the brain of *Piga* mosaic cKO females, we performed western blotting for PIGA in subcortical lysate including the thalamus, basal ganglia, pons, midbrain, and medulla. Western blotting showed a reduction of PIGA protein compared to wild-type littermate controls of approximately 50-75% when normalized to tubulin loading control (Fig.2 C-D). These data confirm PIGA protein reduction in the brain of the mosaic cKO mice.

### CNS GPI deficiency results in decreased survival and weight, but does not grossly affect the structure of the brain

We hypothesized GPI deficiency in the CNS lineage would result in structural defects of the brain including cerebellar hypoplasia, dilated lateral ventricles, and microcephaly as has been observed in a subset of IGD patients [13]. To our surprise, we found no significant structural defects in the cortex of *Piga* mosaic cKO mice at birth (Fig. 3A,B) or up to weaning stages (Fig. 3C,D). We observed mild hypoplasia of the cerebellum in *Piga* mosaic cKO mice as evidenced by H&E staining (Fig. 3E, F). We observed that both the *Piga* mosaic cKO females and the *Piga* hemizygous cKO males gained less weight than their WT littermates and were easily identifiable by this phenotype (Fig. 3G). By P7, all mosaic cKO mutants were approximately half the weight of their WT littermates. We also noticed mutants died postnatally with the oldest mutant only surviving to P23 (Fig. 3H). *Piga* hemizygous cKO males that lack all GPI biosynthesis in the CNS die on average twice as quickly as the mosaic cKO females suggesting a gene dosage effect (Fig. 3H). We hypothesize that the genetic mosaicism in females allows for residual expression of *Piga* allowing longer survive than hemizygous cKO males.

**Figure 3.**
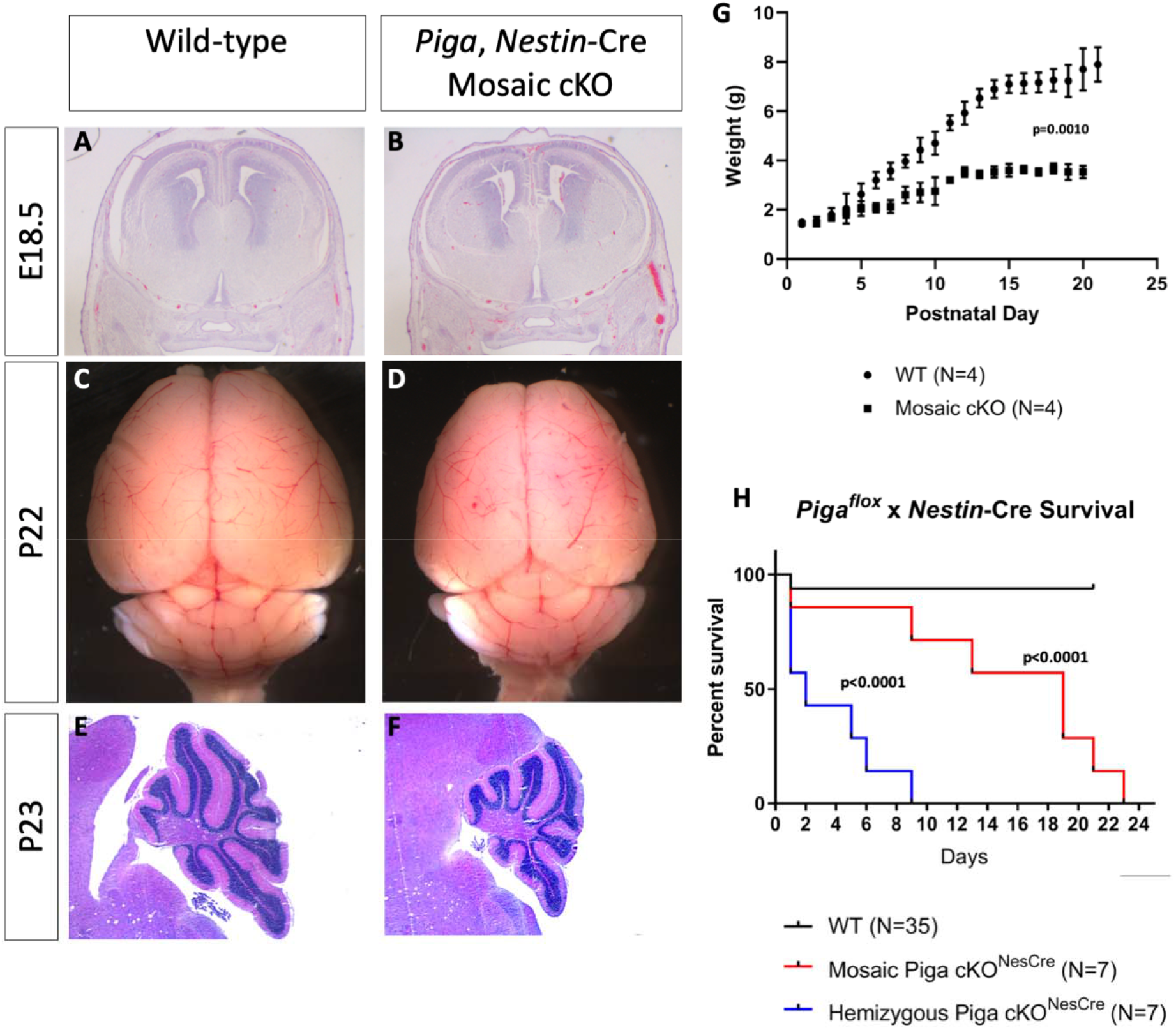
CNS GPI deficiency results in cerebellar hypoplasia, decreased weight gain, and decreased survival. Coronal H&E section of WT and Mosaic cKO cortex at E18.5 (A,B). Gross whole brain of WT and Mosaic cKO at P22 (C,D). H&E section of the cerebellar hemisphere in WT and Mosaic cKO (E,F). Average weight of WT and Mosaic cKO mice from P0 to P15 (G). Survival curve of WT (black), Mosaic cKO (Red), and Hemizygous cKO (Blue) mice (H). All paired images shown at same magnification.

### CNS GPI deficiency results in neurological decline and ataxia

While the cause of death in mutants remains unclear, all mosaic cKO mice developed progressive ataxia, tremor, and a hindlimb clasping phenotype. Moribund mutants were unable to walk but moved their forelimbs and hindlimbs spontaneously while laying on their side in the host cage. Moribund mutants were euthanized at this stage.

Between P10 and P19, we observed a progressive hindlimb clasping phenotype that worsened over time in the mosaic cKO mutants. In the tail suspension test, we scored the hindlimb clasping phenotype according to standard protocols and noted a statistically significantly lower score in the mutants indicating a neurological phenotype (p<0.0001, Fig. 4A-C). Hindlimb clasping is common to many models of neurological disease with a wide variety of underlying pathophysiologies in the brain and spinal cord [23].

**Figure 4.**
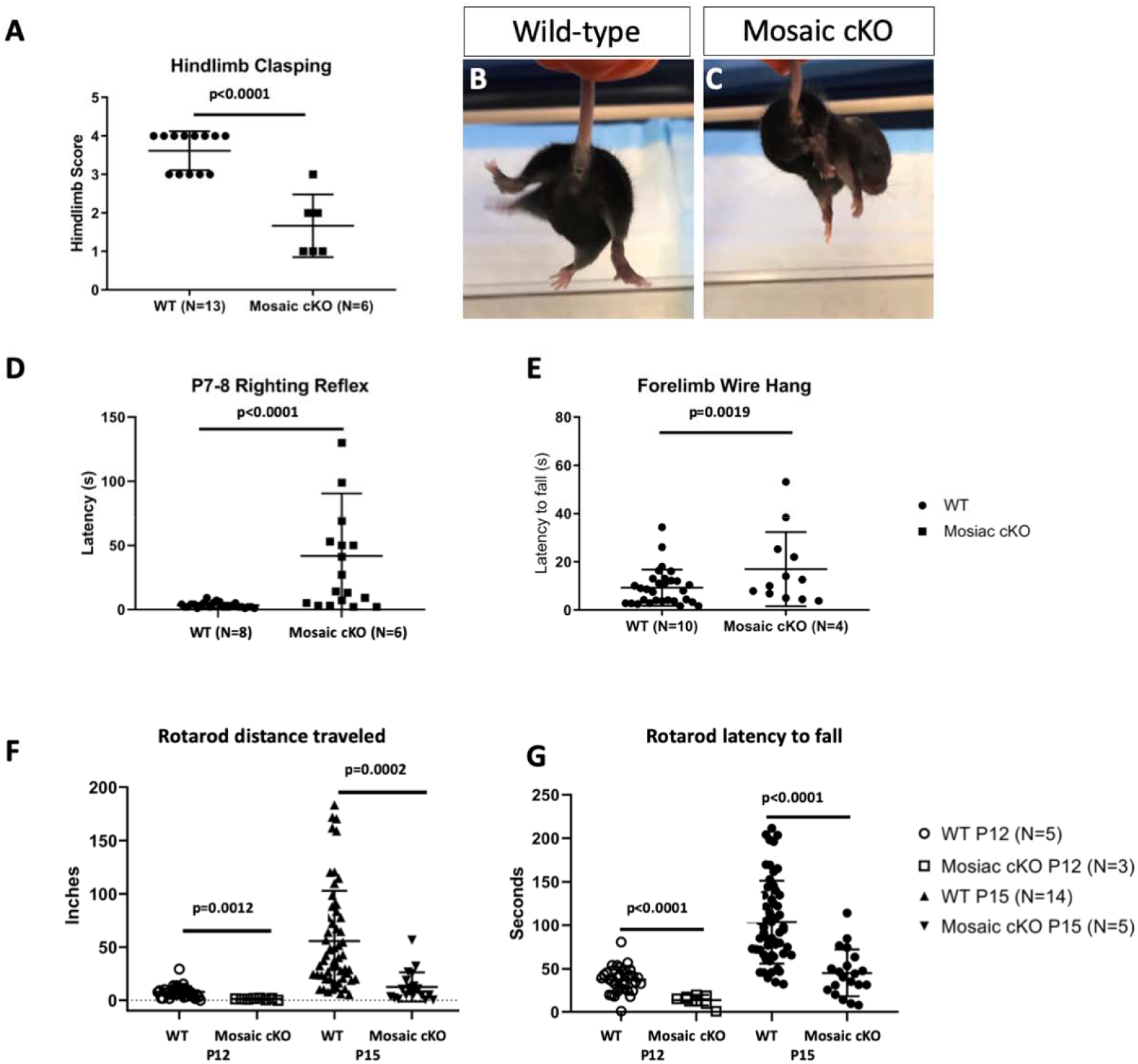
CNS GPI deficiency results in hindlimb clasping, ataxia, and tremor. Hindlimb clasping score of wild-type and mosaic cKO littermates at P16 (A). Images of wild-type and mosaic cKO littermates with mosaic cKO displaying hindlimb clasping (B,C). Surface righting reflex test in wild-type and mosaic cKO littermates (D). Forelimb Wire Hang test in wild-type and mosaic cKO littermates (E). Rotarod testing of wild-type and mosaic cKO littermates at P12 and P15, distance travelled (F) and latency to fall (G).

Given this preliminary indication of neurological disease in postnatal mosaic cKO animals, we sought to test other aspects of neurodevelopment, including developmental landmarks such as reflexes. First, we performed the Righting Reflex test which tests the ability of the mouse to right itself into a prone position after being placed in a supine position. Mosaic cKOs were severely impaired in their ability to right themselves as measured by the latency to right themselves (p<0.0001; Fig. 4D). These data indicate the mutants are defective at either the labryinthe reflex, limb coordination and/or strength as they actively, but unsuccessfully, attempt to right themselves. To test their strength, we performed the forelimb wire hang test in which the mice are suspended on a wire by their forelimbs over a padded surface and measured their latency to fall. Interestingly, we found the mosaic cKOs were able to hang onto the wire statistically longer than their WT littermates, p=0.0019, Fig. 4E). These data indicate the function of the skeletal muscle system and strength is not impaired in the mutant forelimbs. As the mosaic cKOs show deficits in surface righting, but not the forelimb wire hang, we suspected mutants developed defective limb coordination, ataxia.

Over time, the mutants developed ataxia and a persistent tremor (SFig.1). The signs of this are first noticeable at approximately P12 and progressively worsen until approximately P20 when the mutants became moribund. To quantify and further study the ataxia phenotype, we performed rotarod testing. The rotarod assays motor coordination and balance by challenging the mice to maintain their balance on a rotating rod that increases in speed of rotation gradually. We found the mosaic cKOs performed poorly, with significantly decreased travel and latency to fall compared to wild-type littermates at P12 and P15 (ANOVA p=0.0001 for distance traveled and p<0.0001 for latency to fall; Fig. 4F, G). The early lethality of the hemizygous cKO males precluded rotarod analysis. Taken together, all these data from behavioral testing indicate the mosaic cKOs displayed defective motor coordination and balance.

### CNS GPI deficiency delays white matter development

Recent reviews of neural GPI-anchored proteins (GPI-Aps) highlighted the diverse importance of GPI-APs in many neural processes including axon outgrowth, regeneration, synapse formation, neuron cell adhesion, and oligodendrocyte development [24, 25]. We reasoned that failure to properly present one or more specific GPI-APs on the cell membrane may be a cause of the phenotypes observed in the mosaic cKOs. Therefore, we reviewed the phenotypes of mouse knockouts for neural GPI-AP genes in the Mouse Genome Informatics resource. Of the GPI-AP knockout mouse models, our *Piga* mosaic cKO mutants most resemble the phenotype observed in the *contactin 1^-/-^* mouse [26, 27]. *Contactin 1^-/-^* mice weigh significantly less than their wild-type littermates, develop progressive ataxia, and die at approximately P19. Colakoglu et. al. showed *contactin 1* is critical for oligodendrocyte development and mutants lack proper CNS myelination and display defects in cerebellum microorginization [26]. Indeed, multiple mutants in myelin development display an ataxic, tremor, and early death phenotype. Given the phenotypic similarity between *contactin 1^-/-^* mice and *Piga* mosaic cKOs we sought to determine the degree of myelination in *Piga* mosaic cKO mutants.

We performed immunohistochemistry for Myelin Basic Protein (MBP) in wild-type and mosaic cKO mutants at P19 (n=4 for each). We found 3 out of 4 mutants examined showed reduced MBP staining compared to controls indicating that myelination is defective in the mosaic cKO mutants (Fig. 5A-D). We concluded developmental myelination is impaired in *Piga* mosaic cKO mutants. This phenotype has been observed in multiple IGD patients and may be partially responsible for the tremor and early death we observe in the *Piga* mosaic cKO mutants.

**Figure 5.**
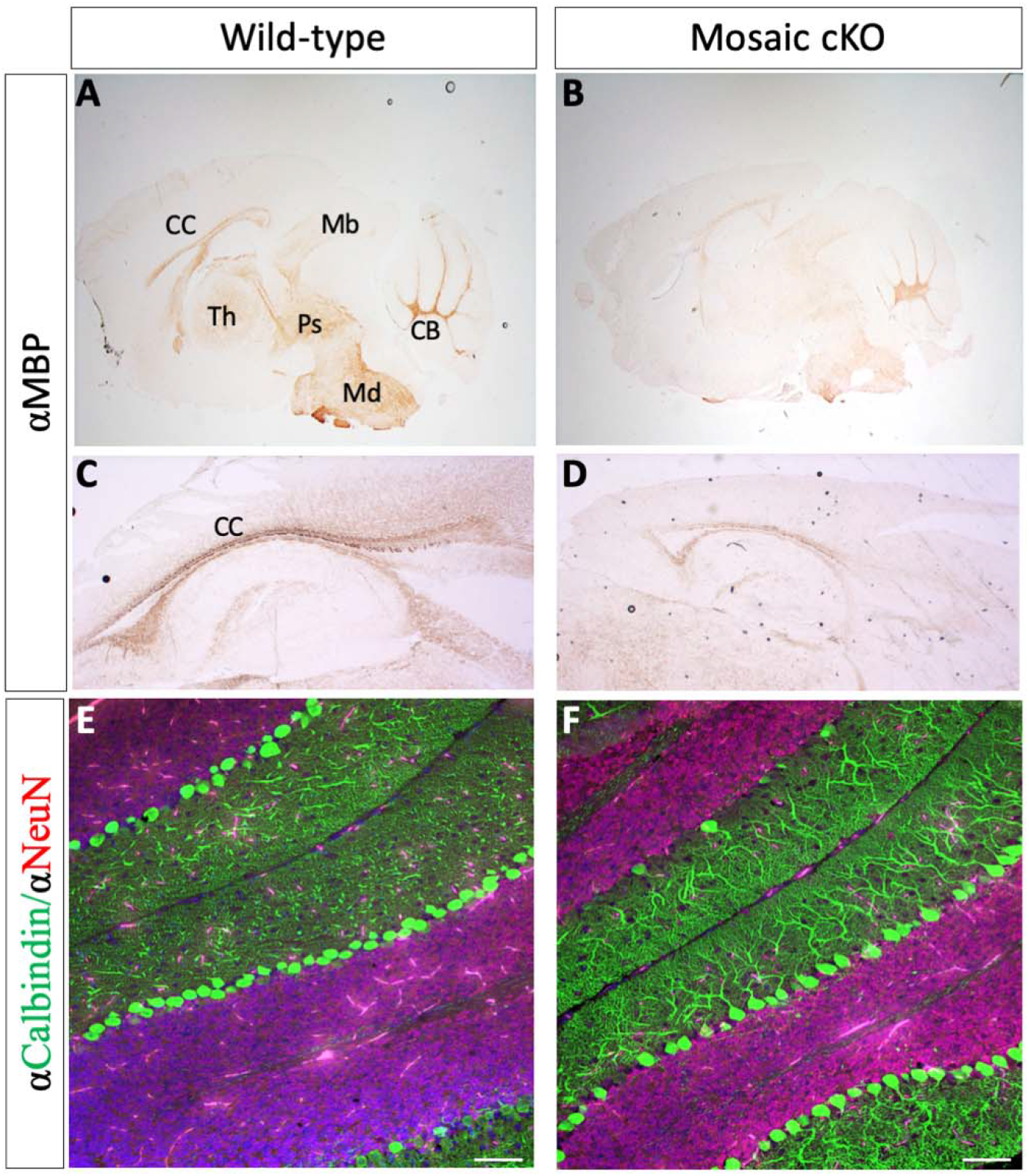
CNS GPI deficiency delays white matter development. Immunohistochemistry of αMBP in wild-type (A, C) and mosaic cKO mice (N=3/4 mutants delayed; B, D). CB= cerebellum, CC=corpus callosum, MB=midbrain, Md=medulla, Ps=pons, Th=thalamus. CNS GPI deficiency impairs Purkinje cell arborization. Purkinje cell (αCalbindin Green) and granule cell (αNeuN Red) immunofluorescence in wild-type and mosaic cKO mice (A, B). Representative images from N= 4/4 Mosaic cKO. (Scale bars = 100 μm. All paired images shown at same magnification.)

### CNS GPI deficiency impairs Purkinje cell dendritic arborization

The ataxia we observed in the *Piga* mosaic cKOs suggested a defect in proper cerebellar function as this is a major center for coordination of motor movement. Defects in either granule cell development or Purkinje cell development can lead to ataxic phenotypes [28–30]. Granule cells are the major excitatory input to Purkinje cells and Purkinje cells are the major inhibitory output to the deep cerebellar nuclei. Given the expression pattern of *Piga* in the postnatal brain, we hypothesized defects in the Purkinje cell layer were responsible for the ataxic phenotype in *Piga* mosaic cKO mice. To assess defects in cerebellar development, we performed immunofluorescence for NeuN to highlight the inner granular layer and Calbindin staining to examine the morphology of the Purkinje cells. The granule cell layer of the cerebellum did not appear to be affected, but the Purkinje cell layer showed marked defects in Purkinje cell dendritic arborization in the molecular layer (n=4/4 mutants examined). We found the dendritic tree to be consistently less elaborated and branched in the *Piga* mosaic cKO mice compared to wild-type littermates (Fig. 5E,F).

To obtain a broader and unbiased understanding of the differences between the wild-type and *Piga* mosaic cKO cerebella, we performed RNA-seq on bulk right cerebellar hemispheres in 3 wild-type and 3 *Piga* mosaic cKO littermates at P20. Differential gene expression analysis identified 176 upregulated genes (Log_2_Fold Change ≥1, P_adjusted_ <0.05), 67 down regulated (Log_2_Fold Change ≤1, P_adjusted_ <0.05) and 17,370 genes that showed no statistical expression difference in the *Piga* mosaic cKO cerebella compared to wildtype controls (Fig. 6A). Of note, the three most highly expressed genes included *Myelin protein zero, Tyrosine hydroxylase,* and *Perlipin 4* (Fig. 6B). Gene Ontology (GO) analysis identified an enrichment for pathways including neuropeptide signaling, iron transport, response to hypoxia, and circadian rhythm for the upregulated genes in the *Piga* mosaic cKO mutant (Fig. 6C).

**Figure 6.**
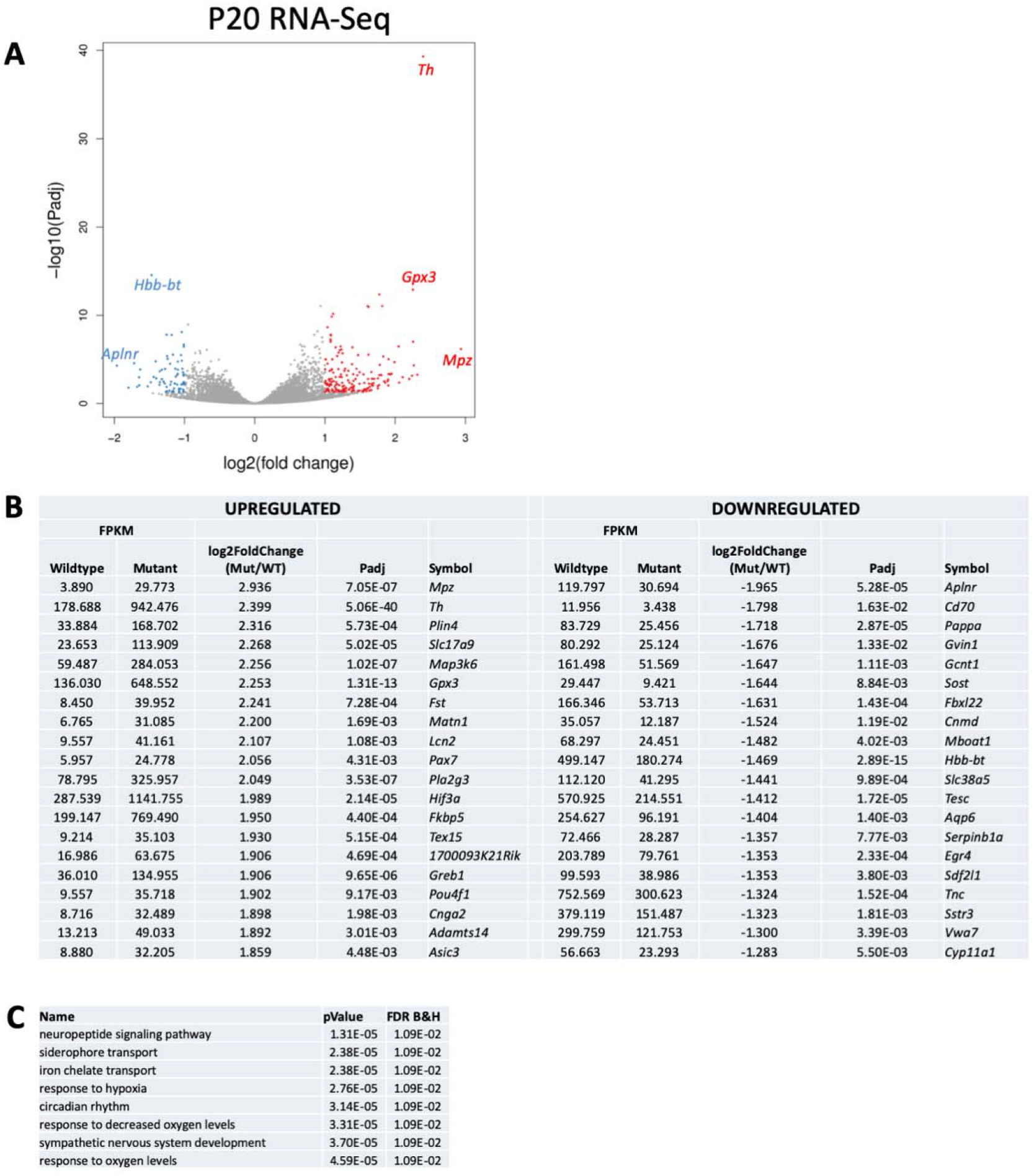
RNA-Seq analysis. (A) Volcano plot of differentially expressed genes from RNA sequencing of wild-type and *Piga* mosaic cKO right cerebellar hemisphere at P20, blue = downregulated genes and red = upregulated genes. (B) Top 20 upregulated and downregulated genes (C). GO analysis of upregulated genes in the *Piga* mosaic cKO cerebella.

## Discussion

In this work we investigated the role of GPI biosynthesis in the developing CNS as it is the most affected organ system in patients with pathogenic variants in GPI biosynthesis genes [31]. We found the initiating enzyme required for GPI biosynthesis, *Piga,* is highly expressed in the developing corpus callosum, periventricular areas of the CNS, and Purkinje cells. We then deleted *Piga* in the CNS/PNS using a conditional approach with *Piga^flox^* mice and *Nestin-Cre* to determine the requirement for GPI biosynthesis in this lineage. We found these mutants do not develop structural defects of the brain as we initially hypothesized. Instead, they developed severe ataxia, tremor, fail to thrive, and died prematurely. As this phenotype resembled other mouse mutants with defects in myelination and IGD patients develop defects in developmental myelination, we sought to determine the degree of myelination in the mutants. We found by αMBP staining mosaic cKO mutants have severe defects in developmental myelination compared to wild-type littermates confirming a requirement for GPI biosynthesis in myelination. We also found severe defects in motor coordination and striking defects in Purkinje cell arborization in the mosaic cKO cerebellum. Similar ataxic phenotypes and reduced lifespan have been observed in other models of CDG including the genetic reductions of *Pmm2, Cog7,* and *Atp6ap2* in *Drosophila* models demonstrating that motor coordination critically requires normal glycosylation through phylogeny [32]. We found gene expression was significantly different in mosaic cKO mice compared to controls with an overexpression of tyrosine hydroxylase, a marker of premature cerebellar development. These data illuminate a novel role for the GPI anchor posttranslational modification in the mouse CNS and provides mechanistic insight into the pathophysiology of IGD.

We were interested to find the expression pattern of *Piga* in the postnatal CNS suggested a role for GPI biosynthesis in white matter development. *Piga* was strongly expressed in the corpus callosum a highly myelinated structure. Myelination is critical to protect axons, increase conduction velocity along the axon, and allow communication between neurons in the CNS especially for motor coordination and learning. Classic models of hypomyelination such as the *rumpshaker* and *quaker* mouse mutants display similar phenotypes as those observed in the mosaic *Piga* cKO mice including truncal ataxia, tremor, and early death [33]. The germline mutants of GPI-anchored contactin family members display a very similar phenotype to that observed in the *Piga* mosaic cKO mice including ataxia, tremor, small body size, and premature death around P20 [26, 27]. Recently, it was shown that *contactin 1* is expressed in oligodendrocytes and is critical for normal myelination [26]. We hypothesize deficiency in the GPI-anchoring of CONTACTIN 1 contributes to the hypomyelination phenotype we observe in the *Piga* mosaic cKO mice. While CONTACTIN1 is a promising candidate, there are several other GPI-APs in the CNS that regulate many related functions including synaptic plasticity, axon outgrowth, and regeneration [24]. We hypothesize that the phenotype observed in the *Piga* mosaic cKO mice highlights the critical early requirements for GPI biosynthesis in the postnatal brain. Indeed, the mosaic cKO mice are moribund by P21 consistent with the lethality seen in a variety of hypomyelination mutants. In contrast, the hemizygous cKO males die even before myelination has really begun, around P10, arguing that GPI biosynthesis is critical for other CNS functions earlier in development. However, we were unable to identify the cause of death in the hemizygous cKO mice and this function of GPI biosynthesis remains unclear.

The most common phenotypes observed in all IGD patients are intellectual disability/developmental delay, and epilepsy. We noticed several episodes of mutants flailing their hindlimbs and forelimbs while on their backs after they fell due to their ataxic gait. They then experienced a sustained period of rigid paralysis with limbs outstretched followed by a short period of inactivity (data not shown). These events may be consistent with a seizure but the early lethality in our mutants precludes a rigorous analysis of the “seizure phenotype” in the mutants by EEG monitoring. These phenotypes are consistent with the “severe” presentation of *Piga* deficiency in which patients die earlier, display severely delayed myelination with thin corpus callosum compared to more “mild” forms of *Piga* deficiency [12, 34]. Further research with a less severe model of GPI biosynthesis deficiency may prove as a better model to study the seizure phenotype as observed in IGD patients. Knock-in of patient variants with the “moderate phenotype” by CRISPR-Cas9 technology, or *Piga* deletion with more restricted Cre transgenes may achieve a more moderate defect in GPI biosynthesis and provide a longer-lived model of IGD with the seizure phenotype.

It has been known for decades that somatic mutations in *Piga* in the hematopoietic lineage result in a hemolytic anemia called Paroxysmal Nocturnal Hemoglobinuria (PNH). PNH red blood cells (RBCs) are missing two critical negative regulators of the complement cascade, the GPI-APs CD55 and CD59. Without GPI biosynthesis, RBCs become the target of aberrant complement activation leading to life threatening hemolytic anemia. Recent research has shown a clinically available complement inhibitor can almost completely halt the PNH disease process [35, 36]. Others have hypothesized the GPI-APs CD59/CD55 may also be critical to protect neurons from complement in the CNS. Indeed, one group derived neurons from *Piga* deficient patient induced pluripotent stem cells and showed that these neurons are more susceptible to complement-mediated lysis in culture [37]. We identified a mild enrichment for immune activation genes in the Piga mosaic cKO cerebellum but they were not among the top hits from GO. The most upregulated genes in mosaic cKO mutants include *myelin protein zero, tyrosine hydroxylase,* and *perilipin 4.* MYELIN PROTEIN ZERO is the major constituent of peripheral myelin in the peripheral nervous system and it is unclear why this gene was overexpressed in our samples. PERILIPIN 4 coats intracellular lipid droplets. *Perilipin 4* overexpression in the mutants may suggest there is storage of lipid droplets in the cells of the mutant. Perhaps defects in GPI biosynthesis result in an accumulation of lipid precursors leading to lipid droplet inclusions in cells.

The strongest signal from our RNA-seq experiment by adjusted P-value was *tyrosine hydroxylase (Th).* Interestingly, though Purkinje cells are not dopaminergic, it has been shown that wildtype Purkinje cells go through a short phase of *Th* expression during development which decreases by P19. In a variety of ataxic mutants including the *pogo, Lrp5/6, β-catenin* cKO, and *dilute* mutants, Purkinje cells abnormally retain *Th* expression to later postnatal stages [38–41]. This retained *Th* expression is thought to mark a delay in Purkinje cell maturation, though the exact role of *Th* in this process is unclear.

Our RNA sequencing results also suggest a role for hypoxia in the pathology of the cerebellar defect observed in the *Piga* Mosaic cKO mice. Hypoxia has been shown to delay the maturation of the cerebellum. How GPI deficiency could result in hypoxia remains unclear. Alterations in lipid metabolism due to the blockade in GPI biosynthesis could allow accumulation of lipid precursors including phosphatidylinositol (PI). If PI were to accumulate then subsequent peroxidation may lead to the generation of reactive oxygen species and a resulting hypoxic response. This remains to be tested. Alternatively, vasculature development may be impaired in the *Piga* mosaic cKO cerebellum leading to regional hypoxia. Further evaluation of vascular development of the *Piga* mosaic cKO would help define this defect.

We propose that our conditional knockout model may serve as an excellent model for preclinical trials of drugs for IGD as the phenotype is robust, quantifiable, and the mosaic cKO survives long enough postnatally to be treated with experimental compounds. The degree of myelination as measured by immunohistochemistry and ataxia as tested with the rotarod serve as convenient end points to examine in experimental settings of preclinical drug trials. A synthetic intermediate of GlcNac-PI would be one promising candidate to test in this model as this intermediate would provide the necessary precursor for GPI biosynthesis missing in *Piga* mutants and possibly rescue GPI expression and the phenotype.

## Materials and Methods

### Animal Husbandry

All animals were maintained through a protocol approved by the Cincinnati Children’s Hospital Medical Center IACUC committee (IACUC2016-0098). Mice were housed in a vivarium with a 12-h light cycle with food and water *ad libitum. Piga^flox^ (B6.129-Piga^tm1^*, #RBRC06211) mice were were previously generated by Taroh Kinoshita and Junji Takeda and obtained from RIKEN [18]. Mice were genotyped for the *Piga^ftox^* allele using Riken’s three primer protocol. *B6.Cg-Tg(Nes-cre)1Kln/J Nestin-Cre mice* (Jackson labs #003771) were genotyped using Jackson lab recommended general Cre genotyping. Progeny of the *Piga^flox^* x Nestin-Cre mice were sacrificed when they became moribund (immobile) at approximately P20. Primers used for genotyping are available in Table S1.

### In Situ Hybridization

Whole E11.5 embryos were fixed overnight in 4% PFA at 4°C and dehydrated through a methanol series. Samples were treated with 4.5μg/mL Proteinase K for 7-13 minutes at room temperature, post-fixed in 4% PFA/0.2% glutaraldehyde and blocked with hybridization buffer prior to hybridization overnight with DIG-labeled in situ probes at 65°C with constant agitation. The samples were washed and incubated with an anti-DIG antibody (Roche #11093274910) o/n at 4°C. Embryos were washed and incubated with BM Purple (Roche #11442074001) from 4 hours at room temperature to o/n at 4°C.

*Piga* plasmid was obtained from Origene (Rockville, MD, #MR222212). Antisense probes were generated from PCR products containing T3 overhangs. A *Piga* antisense probe was generated from 910 base pair PCR product. Primers are listed in Table S1. The PCR products were purified, *in vitro* transcription was performed with digoxigenin-labeled dUTP (Roche #11277073910), and the probe was purified with the MEGAclear Transcription Clean-up kit (Thermo #AM1908) per the manufacturer’s instructions. For sense probes, the plasmids were cut with XhoI restriction enzyme after the coding sequence and T7 RNA polymerase was used for *in vitro* transcription.

### PCR analysis of specific brain regions

Mice were euthanized and tissues were immediately dissected from a variety of brain regions including the cortex, cerebellum, and subcortical areas. Tail biopsies were taken as controls. Tissue was lysed in 50mM sodium hydroxide and boiled on a hot plate for fifteen minutes. The samples were then neutralized with 1M Tris and centrifuged. PCR analysis was performed with 60°C annealing temperature and 34 cycles of amplification for *Piga flox* allele genotyping and a program with 64°C annealing temperature with 34 cycles of amplification for Cre genotyping. Primers are available in Table S1.

### Histology

Brains were dissected and fixed in 10% formalin for 24-48 hours, washed in 70% ethanol, and paraffin embedded by the CCHMC Pathology Core. Brains were sectioned by microtome at 10-20 μm and stained with hematoxylin & eosin using standard methods.

### Weight

Mice used for weight test were weighed individually every day using a standard metric balance from P1 to P21.

### Tail Suspension Test

P15-P18 mice were held by the tail for ten seconds in the air and their hindlimb posture was observed. The mice were given a score from 0-4. 4 indicates the normal hindlimb separation in which they are widely spread when suspended by the tail. Score of 3 means the hindlimbs are more vertical though they barely touch. Score of 2 means the hindlimbs are close and often touching, and 1 indicates profound weakness in which both hindlimbs are almost always clasped together. Score of 0 is given only if the hindlimbs are clasped for the entire ten seconds the mouse is suspended [42].

### Righting Reflex

P7-8 mice were placed supine and the time to flip over onto their abdomens was measured in seconds and recorded as a latency to right themselves to a prone position [42]. Three trials were performed consecutively and each trial time was plotted.

### Forelimb Wire Hang

Mice were placed on a thin wire suspended by their forelimbs over a padded drop zone. The pups were placed on the wire such that the experimenter could observe their grip on the wire. The pups were then released and the time before they fell off the wire was measured in seconds and recorded as a latency to fall [42]. Three trials were performed in sequence and each trial latency to fall was plotted.

### Rotarod

Rotarod was performed by the CCHMC Animal Behavior Core [43]. Researchers were blinded to the genotypes of the animals tested. Rotarod apparatus (San Diego Instruments) was used with SDI software with the 4 to 40 RPM (revolutions per minute) protocol for mice. The apparatus has four chambers and four mice were tested at the same time. The photobeam sensor detects and records the time and distance each mouse travels before falling from the rod. The test was allowed to run for 360 seconds. The test starts at 4RPM for 30 seconds and increases to 16RPM for 110 seconds, then 28RPM for 110 seconds, and finally 40RPM for 110 seconds. The latency to fall in seconds and distance traveled was recorded automatically and the mice were allowed to rest for 15 minutes before each successive trial. Four trials were performed sequentially on each mouse.

### Immunohistochemistry

Immunohistochemistry was performed on formalin-fixed, paraffin embedded brain tissue harvested from P19 animals. Briefly, tissue was sectioned at 10-20μm, sections were blocked for one hour at room temperature in 4% normal goat serum in PBST, and incubated in primary antibody (1:500 chicken antimouse Myelin Basic Protein antibody, Aves Inc, #MBP) overnight at 4°C. The next day, slides were washed in PBS, and incubated in 1:500 biotinylated goat anti-chicken antibody (Aves Inc, #B-1005) for one hour at room temperature. The slides were washed and incubated in ABC mix (Vectastain ABC HRP Kit, #PK-4000) for 1 hour at room temperature. The slides were washed and developed in 0.5 mg/mL DAB (Sigma) activated with 30% hydrogen peroxide. Slides were incubated in DAB for approximately 5 minutes, washed in PBS, sealed with Cytoseal, and imaged by light microscopy.

### Immunofluorescence

P0-P23 mice were euthanized with isoflurane and cervical dislocation and their brains were microdissected in PBS. They were fixed in 4% PFA o/n, equilibrated in 30% sucrose o/n, cryo-embedded in OCT, and sectioned from 10-20μM with cryostat. Antigen retrieval was performed with citrate retrieval buffer, blocked in 4% normal goat serum, incubated in primary antibodies o/n at 4°C: rabbit anti-PIGA (Proteintech #13679-1-AP, 1:200), mouse anti-NeuN (Millipore #MAB377, 1:1,000), Rat anti-CD68 (Biorad #MCA1957T, 1:1,000), rabbit anti-Calbindin (abcam #ab25085, 1:4,000), and Rat anti-Mylein Basic Protein (Aves #MBP, 1:500). Sections were incubated with Alexafluor 488-congugated goat antirabbit (Thermo #A11008, 1:1,000) and Alexafluor 594 conjugated goat anti-mouse (Thermo #A11008, 1:1,000) secondary antibodies and counterstained with DAPI. Sections were imaged on Nikon C2 confocal 703 microscope.

### Western Immunoblotting

Brains were microdissected and the cortex and cerebellum were isolated. Subcortical tissue was lysed in 800μL RIPA buffer+ Protease inhibitor. Lysate protein concentration was determined by BCA assay and electrophoresis was performed on a 10% Tris-glycine gel. Protein was transferred to a PVDF membrane, blocked in Odyssey blocking buffer and incubated o/n at 4°C with Rabbit anti-PIGA (Proteintech #13679-1-AP, 1:1,000) and Mouse anti-Tubulin (Sigma #T6199, 1:1,000) antibodies. Membranes were washed and incubated for 1 hour in goat anti-rabbit IRDye 800CW (LICOR # 926-32211, 1:15,000)and goat antimouse IRDye 680Rd (LICOR, #926-68070, 1:15,000) antibodies and visualized on a LICOR Odyssey imaging system. Relative protein concentration was determined by normalizing PIGA signal to Tubulin signal in Image Studio Lite Ver 5.2.

### RNA Sequencing

3 WT and 3 mosaic cKO cerebella were bisected along the midline of the cerebellar vermis and the right half was snap frozen on dry ice. RNA was isolated and pooled samples of each genotype were used for paired end bulk RNA sequencing (BGI Americas, Cambridge, MA). mRNA molecules were purified from total RNA using oligo(dT) attached magnetic beads. mRNA molecules were fragmented into small pieces using fragmentation reagent and first strand cDNA was generated using random hexamer primed reverse transcription, followed by a second strand cDNA synthesis. The synthesized cDNA was subjected to end repair and 3’ adenylated. Adapters were ligated to the ends of these 3’ adenylated cDNA fragments. PCR was used to amplify the cDNA fragments with adapters from previous step. PCR products were purified with Ampure XP Beads (AGENCOURT) and dissolved in EB solution. Library was validated on the Agilent Technologies 2100 bioanalyzer. The double stranded PCR products were heat denatured and circularized by the splint oligo sequence. The single strand circle DNA (ssCir-DNA) were formatted as the final library. The library was amplified with phi29 to make the DNA nanoball (DNB) which had more than 300 copies of one molecule. The DNBs were load into the patterned nanoarray and single end 50(pair end 100/150) bases reads were generated in the way of combinatorial Probe Anchor Synthesis (cPAS). Analysis was performed by BGI RNA Sequencing services which includes a proprietary analysis pipeline.

### Statistical Analysis

Statistical analysis was performed using Graphpad Prism (GraphPad Software, San Diego, CA). Survival analysis was performed with a log-rank (Mantel-Cox) test. Rotarod analysis was performed with a oneway ANOVA followed by unpaired t-tests. All other tests were unpaired t-tests.

---

**Table S1.**
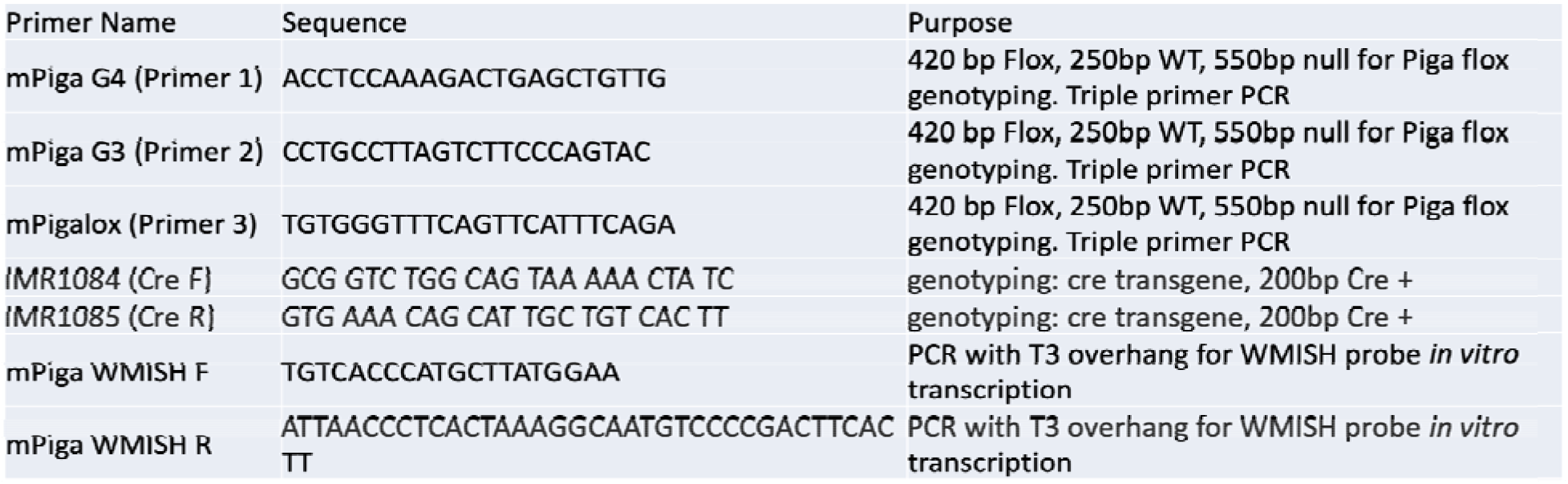
PCR Primers

## References

1. Kinoshita, T., Biosynthesis and deficiencies of glycosylphosphatidylinositol. Proc Jpn Acad Ser B Phys Biol Sci, 2014. 90(4): p. 130–43.

2. Bellai-Dussault, K., et al., Clinical variability in inherited glycosylphosphatidylinositol deficiency disorders. Clinical Genetics, 2019. 95(1): p. 112–121.

3. Kinoshita, T., Glycosylphosphatidylinositol (GPI) Anchors: Biochemistry and Cell Biology: Introduction to a Thematic Review Series. Journal of Lipid Research, 2016. 57(1): p. 4–5.

4. Freeze, H.H., Understanding Human Glycosylation Disorders: Biochemistry Leads the Charge. Journal of Biological Chemistry, 2013. 288(10): p. 6936–6945.

5. Knaus, A., et al., Characterization of glycosylphosphatidylinositol biosynthesis defects by clinical features, flow cytometry, and automated image analysis. Genome Medicine, 2018.10.

6. Knaus, A., et al., Rare Noncoding Mutations Extend the Mutational Spectrum in the PGAP3 Subtype of Hyperphosphatasia with Mental Retardation Syndrome. Human Mutation, 2016. 37(8): p. 737–744.

7. Pagnamenta, A.T., et al., Analysis of exome data for 4293 trios suggests GPI-anchor biogenesis defects are a rare cause of developmental disorders. European Journal of Human Genetics, 2017. 25(6): p. 669–679.

8. Nguyen, T.T.M., et al., Mutations in PIGS, Encoding a GPI Transamidase, Cause a Neurological Syndrome Ranging from Fetal Akinesia to Epileptic Encephalopathy. American Journal of Human Genetics, 2018. 103(4): p. 602–611.

9. Swoboda, K.J., et al., A Novel Germline PIGA Mutation in Ferro-Cerebro-Cutaneous Syndrome: A Neurodegenerative X-Linked Epileptic Encephalopathy With Systemic Iron-Overload. American Journal of Medical Genetics Part A, 2014. 164(1): p. 17–28.

10. Kuki, I., et al., Vitamin B-6-Responsive Epilepsy Due to Inherited Gpi Deficiency. Neurology, 2013. 81(16): p. 1467–1469.

11. Thompson, M.D., et al., Hyperphosphatasia with neurologic deficit: A pyridoxine-responsive seizure disorder? Pediatric Neurology, 2006. 34(4): p. 303–307.

12. Tarailo-Graovac, M., et al., The genotypic and phenotypic spectrum of PIGA deficiency. Orphanet Journal of Rare Diseases, 2015. 10.

13. Johnston, J.J., et al., The Phenotype of a Germline Mutation in PIGA: The Gene Somatically Mutated in Paroxysmal Nocturnal Hemoglobinuria. American Journal of Human Genetics, 2012. 90(2): p. 295–300.

14. Lukacs, M., et al., Glycosylphosphatidylinositol biosynthesis and remodeling are required for neural tube closure, heart development, and cranial neural crest cell survival. Elife, 2019. 8.

15. Ahrens, M.J., et al., Convergent extension movements in growth plate chondrocytes require gpi-anchored cell surface proteins. Development, 2009. 136(20): p. 3463–3474.

16. Tarutani, M., et al., Tissue-specific knockout of the mouse Pig-a gene reveals important roles for GPI-anchored proteins in skin development. Proceedings of the National Academy of Sciences of the United States of America, 1997. 94(14): p. 7400–7405.

17. Visconte, V., et al., Phenotypic and functional characterization of a mouse model of targeted Pig-a deletion in hematopoietic cells. Haematologica-the Hematology Journal, 2010. 95(2): p. 214–223.

18. Nozaki, M., et al., Developmental abnormalities of glycosylphosphatidylinositol-anchor-deficient embryos revealed by Cre/loxP system. Laboratory Investigation, 1999. 79(3): p. 293–299.

19. Tronche, F., et al., Disruption of the glucocorticoid receptor gene in the nervous system results in reduced anxiety. Nature Genetics, 1999. 23(1): p. 99–103.

20. Zhang, Y., et al., An RNA-Sequencing Transcriptome and Splicing Database of Glia, Neurons, and Vascular Cells of the Cerebral Cortex (vol 35, pg 11929, 2014). Journal of Neuroscience, 2015. 35(2): p. 864–866.

21. Giusti, S.A., et al., Behavioral phenotyping of Nestin-Cre mice: Implications for genetic mouse Models of psychiatric disorders. Journal of Psychiatric Research, 2014. 55: p. 87–95.

22. Graus-Porta, D., et al., beta 1-class integrins regulate the development of laminae and folia in the cerebral and cerebellar cortex. Neuron, 2001. 31(3): p. 367–379.

23. Lalonde, R. and C. Strazielle, Brain regions and genes affecting limb-clasping responses. Brain Research Reviews, 2011. 67(1-2): p. 252–259.

24. Um, J.W. and J. Ko, Neural Glycosylphosphatidylinositol-Anchored Proteins in Synaptic Specification. Trends in Cell Biology, 2017. 27(12): p. 931–945.

25. Freeze, H.H., et al., Neurological Aspects of Human Glycosylation Disorders. Annual Review of Neuroscience, Vol 38, 2015. 38: p. 105-+.

26. Colakoglu, G., et al., Contactin-1 regulates myelination and nodal/paranodal domain organization in the central nervous system. Proceedings of the National Academy of Sciences of the United States of America, 2014. 111(3): p. E394–E403.

27. Berglund, E.O., et al., Ataxia and abnormal cerebellar microorganization in mice with ablated contactin gene expression. Neuron, 1999. 24(3): p. 739–750.

28. Cendelin, J., From mice to men: lessons from mutant ataxic mice. Cerebellum Ataxias, 2014. 1: p. 4.

29. Wang, J.Y., et al., Sun1 deficiency leads to cerebellar ataxia in mice. Disease Models & Mechanisms, 2015. 8(8): p. 957-+.

30. Becker, E.B.E., et al., A point mutation in TRPC3 causes abnormal Purkinje cell development and cerebellar ataxia in moonwalker mice. Proceedings of the National Academy of Sciences of the United States of America, 2009.106(16): p. 6706–6711.

31. Bellai-Dussault, K., et al., Clinical variability in inherited glycosylphosphatidylinositol deficiency disorders. Clin Genet, 2018.

32. Frappaolo, A., et al., Modeling Congenital Disorders of N-Linked Glycoprotein Glycosylation in Drosophila melanogaster. Frontiers in Genetics, 2018. 9.

33. Duncan, I.D. and A.B. Radcliff, Inherited and acguired disorders of myelin: The underlying myelin pathology. Exp Neurol, 2016. 283(Pt B): p. 452–75.

34. Kato, M., et al., PIGA mutations cause early-onset epileptic encephalopathies and distinctive features. Neurology, 2014. 82(18): p. 1587–96.

35. Kinoshita, T., Molecular genetics, biochemistry, and biology of PNH. Rinsho Ketsueki, 2017. 58(4): p. 353–362.

36. Hillmen, p., et al., The complement inhibitor eculizumab in paroxysmal nocturnal hemoglobinuria. New England Journal of Medicine, 2006. 355(12): p. 1233–1243.

37. Yuan, X., et al., A hypomorphic PIGA gene mutation causes severe defects in neuron development and susceptibility to complement-mediated toxicity in a human iPSC model. Plos One, 2017. 12(4).

38. Jeong, Y.G., M.K. Kim, and R. Hawkes, Ectopic expression of tyrosine hydroxylase in Zebrin II immunoreactive Purkinje cells in the cerebellum of the ataxic mutant mouse, pogo. Brain Res Dev Brain Res, 2001. 129(2): p. 201–9.

39. Huang, Y., et al., Lrp5/6 are reguired for cerebellar development and for suppressing TH expression in Purkinje cells via beta-catenin. Mol Brain, 2016. 9: p. 7.

40. Sawada, K., et al., Abnormal expression of tyrosine hydroxylase immunoreactivity in Purkinje cells precedes the onset of ataxia in dilute-lethal mice. Brain Res, 1999. 844(1-2): p. 188–91.

41. Sawada, K., et al., Abnormal expression of tyrosine hydroxylase immunoreactivity in cerebellar cortex of ataxic mutant mice. Brain Res, 1999.829(1-2): p. 107–12.

42. Feather-Schussler, D.N. and T.S. Ferguson, A Battery of Motor Tests in a Neonatal Mouse Model of Cerebral Palsy. Jove-Journal of Visualized Experiments, 2016(117).

43. McAuliffe, J.J., et al., Desflurane, Isoflurane, and Sevoflurane Provide Limited Neuroprotection against Neonatal Hypoxia-Ischemia in a Delayed Preconditioning Paradigm. Anesthesiology, 2009. 111(3): p. 533–546.

